# Evolutionary trade-offs between functional and immune selection shape multigene families in pathogens

**DOI:** 10.1101/2025.06.24.661038

**Authors:** Qi Zhan, Mercedes Pascual, Qixin He

**Author notes:** Correspondence should be addressed to Qixin He.

## Abstract

Major surface antigens in many pathogens are encoded by rapidly diversifying multigene families, generating fitness variation through antigenic and functional differences. These variations align with the niche and absolute fitness axes of Modern Coexistence Theory (MCT). Yet, how such gene families evolve along these axes under competition for hosts and across transmission gradients remains poorly understood, as prior MCT studies have not explicitly accounted for evolutionary dynamics in high dimensions. We use a stochastic computational model of *Plasmodium falciparum* transmission to examine how transmission intensity and selection shape *var* multigene family evolution and composition within parasite genomes. Results show that selection alone cannot maintain the observed stable ratio of two gene groups within parasite genomes, indicating that group-based classifications do not clearly reflect transmission strategy or virulence. When a trade-off exists between diversification rates and absolute fitness, strong immune selection under high transmission favors fast-recombining genes while attenuating functional selection on *R*_0_-associated traits. In general, stronger immune selection increases the invasion probability of novel antigens and the niche differentiation among parasite genomes, while reducing the variance in gene-level transmissibility and expression duration, and therefore *R*_0_. This outcome, combining enhanced niche differentiation and reduced absolute fitness variation, departs from MCT predictions.

## Background

The evolutionary arms race between infectious pathogens and their hosts reflects the constant struggle of pathogens to proliferate and the need for infected hosts to curb their growth^1,2^ . Multigene families play key roles in the adaptation of both sides. The major histocompatibility complex (MHC) genes in vertebrates encode highly polymorphic cell surface proteins that are essential for adaptive immune responses^3^ . Similarly, many pathogens across protozoa, bacteria, and fungi employ multigene families to encode diverse surface antigens, which can undergo rapid innovation, generating extensive antigenic variation at both within-host and population levels to facilitate immune evasion^4^ . Although evolutionary models for MHC have been extensively explored^5,6,7^, the birth-death processes of individual member genes within antigen-encoding multigene families of parasite genomes, have been less studied. Understanding the evolution of these member genes and the associated traits of their strains, as well as the drivers of this evolution, is crucial to predict changes in the ability of an important class of pathogens to cause severe disease and to identify strategies to reduce the disease burden in human and livestocks^8^ . Within theoretical ecology, modern coexistence theory (MCT) for competitive systems helps explain how species interactions shape ecological communities. It does so by organizing trait variation along two major axes. The first axis concerns traits that influence competitive ability and consequently fitness in an “absolute” manner, independent of their frequencies, whose differences typically oppose coexistence under conditions of high competition. Traits of this kind are related to demography and intrinsic growth rates. The second axis concerns traits whose competitive effects are purely frequency-dependent, whose differences have a “stabilizing” impact, promoting coexistence through niche differentiation and resource partitioning^9^ . Coexistence is favored by “equalizing” differences in absolute fitness and by enhancing niche differences. This framework has recently been extended to host-pathogen systems^10^ . In these previous studies, including the recent extension specific to pathogens, the focus has been on pairwise interactions and therefore on low-diversity systems, with mathematical formulations that rely on preexisting diversity but do not consider the emergence of novelty and therefore explicit evolutionary dynamics. Extending MCT to high-dimensional systems and phenotypes composed of multiple explicit traits, with ecological and evolutionary processes occurring on similar timescales, remains a major challenge^11^ . Nevertheless, this conceptual framework allows us to categorize here the trait variation among individual gene members of a given multigene family along these same two axes: one for absolute fitness differences (basically the basic reproductive number *R*_0_ of a gene), related to virulence, transmissibility, and duration of expression, and another for niche differences due to antigenic variation and associated immune evasion. For the latter, we do consider here the rate of novel gene generation which facilitates such immune escape. We can then ask how variation along these axes influences the persistence and turnover of individual member genes, and therefore, the coexistence and composition of gene properties, within parasite genomes and populations.

The human malaria parasite *Plasmodium falciparum* provides a good model system for addressing how both functional selection for high absolute fitness and immune selection for antigenic novelty shape the coexistence of individual member genes within a multigene family. It is a major human pathogen whose extensive within-host and population-level antigenic diversity underlies resilience against intervention efforts at high transmission^12,13,14,15^ . The *var* multigene family encodes the *Plasmodium falciparum* erythrocyte membrane protein 1 (*Pf* EMP1), the major variant surface antigen of the blood-stage of infection. From this perspective, strain variation occurs within a high-dimensional trait space, with evolution influenced by the population dynamics of the disease, including the competitive interactions of strains for hosts as a function of their immune memory. Evolutionary change occurs in the frequency of both genes and genotypes within the population and involves the emergence of new genes through mitotic recombination, immigration, and mutation, as well as the generation of new strains through meiotic recombination.

More specifically, each parasite carries around 50-60 *var* genes, each encoding a distinct antigenic form of *Pf* EMP1^16^ . Typically, only one or a few genes are expressed at a time, enabling sequential switching of antigenically distinct surface antigens^17,18^ . *var* genes and their corresponding *Pf* EMP1 proteins exhibit trait variation along the two above-mentioned axes^19,20,21,22^ . Specifically, functionally (or with respect to the axis of absolute fitness), *Pf* EMP1 binds to host endothelial receptors, preventing parasites from splenic clearance. Thus, a higher binding affinity confers a higher absolute fitness of parasites^19^, influencing the virulence of the pathogen^23^, the transmissibility and the duration of expression, or their combined effect measured by *R*_0_. Regarding the frequency-dependent axis, *var* genes are known to exhibit different rates of mitotic recombination, generating novel offspring genes encoding new surface proteins that improve immune evasion^24^ . The major groups of *var* genes, upsA and upsB/C, are classified by their 5’-flanking regions (ups), which regulate gene expression^16^ . These two groups are believed to occupy distinct regions of the trait space along these two axes, reflecting diversification adapted to the varying conditions of transmission and the host immune response^25^ . UpsA genes are thought to encode highly virulent *Pf* EMP1s, which underlie severe disease outcomes. In contrast, upsB/C genes are considered to have evolved to facilitate fast mitotic recombination for immune evasion, potentially compromising their ability to effectively bind to host endothelial receptors. An important observation is that repertoires of genes (individual parasite genomes) and isolates (sets of individual parasite genomes co-infecting individual hosts) in the field consistently show an approximately constant proportion of the two ups groups across different geographical locations and transmission intensity, about 20% upsA and 80% upsB/C ^16,12,14,26,27^ . The mechanism underlying this stable proportion remains unclear.

Using a stochastic, computational agent-based model of the transmission system, we examine here the eco-evolutionary dynamics of the *var* multigene family to address how transmission intensity and associated immune selection shape the composition of multigene families within individual parasite genomes. We ask under what conditions pathogens evolve to optimize virulence, transmissibility, and duration of expression, leading to the dominance of genes with optimal trait values within genomes. Specifically, we investigate how trade-offs among gene-level traits influence the birth-death processes of individual member genes within multigene families along a transmission gradient.

Although trade-offs shaping pathogen evolution have been extensively studied^28,29^, existing theoretical work has largely focused on single-locus systems. Our study is the first to explicitly examine how trade-offs shape the epidemiological and evolutionary dynamics of highly diverse multigene systems. In particular, we ask whether trade-offs between the two axes reflecting different transmission strategies can promote the stable coexistence of genes adapted to each strategy within parasite genomes and at different transmission intensities. We specifically consider four different scenarios with an extension of the stochastic agent-based model (ABM) in Zhan et al.^15^ . The first scenario allows traits to vary freely along both axes without any trade-off constraint. The second and third ones introduce variation along each axis and trade-offs within, but not between, the two axes. These consist of respective associations between transmissibility and duration of expression within the axis for absolute fitness differences, and between innovation rate and load of novelty within the axis for niche differences. The final scenario allows variations along both axes while introducing a trade-off between them. We find that no scenario can generate the emergence of stable proportions of gene groups. Furthermore, high transmission rates generate higher invasion probabilities of novel antigens and high diversity at both the gene and genome levels, with parasite genomes exhibiting low overlap in gene composition and correspondingly large niche differences. However, high transmission also selects for genes with high mitotic recombination rates and leads to the homogenization of functional differences among genes. This “equalizing” of functional differences with reduced variance across genes at high niche differences, opposes the expectation of MCT for purely ecological dynamics. Understanding the evolutionary outcomes of these trade-offs helps generate hypotheses for the maintenance of the empirically observed, approximately stable ratio of the two ups groups within individual parasite genomes, clarifies the relationship between ups groups and transmission strategies, and has important implications for identifying pathologically and clinically relevant subsets of *var* genes based solely on sequence information–an ongoing challenge given their hyperdiversity.

## Methods

### Trait variation, trade-offs, and ups groups of *var* genes

#### *R*_0_ of *var* genes

The basic reproductive number, *R*_0_, is defined as the expected number of secondary infections generated by a single infected case in a fully susceptible population in the absence of interventions. *R*_0_ is typically defined at the genotype or strain level. In this study, however, we do not directly specify or parameterize strain-level *R*_0_. Instead, we specify the underlying biological parameters that govern gene-specific traits, along with a transmission parameter that contributes to transmission intensity (see the next paragraph for details). Together, these parameters determine the reproductive potential of a strain through the sequential expression of genes within its multigene family. We emphasize that the reproductive potentials of genes and strains are intrinsically intertwined. The effective reproductive number of a gene depends not only on its own traits, but also on the strain in which it is embedded and on the transmission and expression dynamics of other genes within that strain, since transmission occurs at the level of the strain as a whole.

At the gene level–i.e., while a particular gene is actively expressed (see the section on Within-host dynamics in the supplementary material for details)–the transmission success of that gene, and its contribution to the overall transmission success of the strain in which it resides, depends on three components: the population-level host contact rate (*b*), gene-specific transmissibility (*T*), and the gene-specific duration of expression (*d*). Gene-specific transmissibility (*T*) is defined as the probability that a parasite strain expressing a given *var* gene–under single infection and in the absence of co-infecting strains–is successfully transmitted to a susceptible host during a transmission event. The duration of expression (*d*) is described in detail in the section on Within-host dynamics in the supplementary material. Together, *T* and *d* determine the gene-level contribution to transmission success, independent of the population-level host contact rate.

Whereas *T* and *d* are intrinsic properties of individual genes, the host contact rate *b* is specified at the host-population level and is shared across all strains and genes. Accordingly, when visualizing variance in trait values, we focus on the product *T × d*, which captures the gene-level component of transmission success independently of host contact rates.

### Two different selection pressures from hosts and the resulting transmission strategies of *var* genes

*Pf*EMP1s, encoded by *var* genes, are transported to and expressed on the surface of infected erythrocytes (IEs) (supplementary figure S1a). These *Pf*EMP1s are responsible for IE adhesion, including cytoadherence to endothelial cells (often referred to simply as cytoadherence or cytoadhesion) and rosetting with uninfected erythrocytes, which prevent splenic clearance and enhance parasite reproduction rates^19^ . Furthermore, *Pf*EMP1s are the major blood-stage antigens that mediate IEs’ binding to various immune system cells, leading to significant immunological consequences^19^ . Thus, *var* genes and more generally many other infectious pathogen antigen-encoding genes^30^ experience two different selection pressures from hosts during infections. On the one hand, gene variants with enhanced cytoadhesion allow parasites carrying a higher proportion of these variants to grow faster and more efficiently in naive hosts, increasing transmissibility, prolonging duration of expression, and ultimately increasing the basic reproductive number (*R*_0_). This generates positive, directional selection favoring genes with optimal absolute fitness (*R*_0_) regardless of their frequency. We refer to this process as functional selection acting along the axis of absolute fitness differences. On the other hand, gene variants with novel epitopes, unrecognized by preexisting antibody responses, confer a within-host growth advantage in immune hosts, favoring parasites that carry a higher proportion of these variants. This generates negative frequency-dependent selection, whereby common genes incur a fitness disadvantage due to widespread immune memory, while rare genes are favored through immune evasion. Because this frequency-dependent selection promotes coexistence via niche differentiation and resource partitioning, we refer to it as immune selection acting along the axis of niche differences. Consequently, *var* genes can adopt one or both of two distinct transmission strategies: (1) an intrinsic growth advantage within naive hosts, leading to higher virulence, transmissibility, or longer duration of expression–collectively yielding a higher *R*_0_ (the y-axis in the supplementary figure S1b); (2) rapid innovation that generates novel genes capable of evading host immunity, thereby expanding the pool of susceptible hosts through niche differentiation and resource partitioning (the x-axis in the supplementary figure S1b, shown there as niche overlap, the complement of niche differences).

### Potential trade-offs between traits of *var* genes

Trade-offs^31^ may operate along each of the two axes individually, as well as between them, as parasites adopt one or both transmission strategies. Given the limited understanding of genotype-phenotype mapping, we consider four scenarios (Table 1): (1) genes vary freely along both axes and evolve without trade-off constraints; (2) genes vary only along the axis of absolute fitness differences and evolve under trade-offs among traits associated with this axis; (3) genes vary only along the axis of niche differences and evolve under trade-offs between traits associated with this axis; and (4) genes vary along both axes and evolve under trade-offs between traits associated with the two axes. This final scenario captures the tug-of-war between functional selection and immune selection.

**Table 1:** The four scenarios considered. G1 refers to group 1 genes and G2 refers to group 2 genes.

|  | Scenario (1) | Scenario (2) | Scenario (3) | Scenario (4) |
| --- | --- | --- | --- | --- |
| Transmissibility (Trans) | $G1 = 2 \times G2$ | $G1 = 2 \times G2$ | $G1 = G2$ | $G1 = 2 \times G2$ |
| Duration (Dur) | $G1 = \frac{4}{3} \times G2$ | $G1 = \frac{12}{13}$ (or $\frac{1}{2}$) $\times G2$ | $G1 = G2$ | $G1 = \frac{4}{3} \times G2$ |
| Trans × Dur | $G1 > G2$ | $G1 > \text{(or } = \text{)} G2$ | $G1 = G2$ | $G1 > G2$ |
| Recombination Rate | $G1 \ \& \ G2 \in \text{(Low, High)}$ | $G1 = G2 = 0$ | $G1 < G2$ | $G1 < G2$ |
| Recombination Load | $G1 = G2 = 0$ | Not Applicable | $G1 < G2$ | $G1 < G2$ |

The absolute fitness of genes and their associated pathogens is determined by the combined effects of multiple traits, including transmissibility and duration of expression. In contrast, niche differentiation and resource partitioning are quantified by gene innovation rates, driven primarily by mitotic recombination. In scenario (1), newly created genes may vary freely in transmissibility (high or low), duration of expression (long or short), and thus *R*_0_, as well as in mitotic recombination rate (fast or slow). Genes with higher transmissibility may be associated with higher parasitemia, eliciting stronger immune response in infected hosts and consequently shortening their duration of expression. This generates a trade-off between transmissibility and duration of expression whose strength may vary^32,33,34,35^, ranging from weak (partial compensation) to strong (full compensation), with the latter equalizing absolute fitness (*R*_0_) across genes. In scenario (2), we consider both cases. Under a strong trade-off, an effectively neutral regime emerges in which no deterministic competitive winner exists, allowing genes with different transmissibility–duration combination to coexist for extended periods before stochastic drift drives some to extinction.

Mitotic recombination can generate functional, novel offspring genes that no longer compete with existing genes for the same immune space. However, recombination also incurs a cost–termed recombination load–through the production of nonfunctional offspring that replace their parental genes and are cleared immediately, thereby reducing infection duration and strain fitness. In scenario (3), we examine whether this recombination load undermines selection for fast mitotic recombination. As gene variants, particularly immunodominant regions, diversify, binding affinities may deteriorate, introducing a trade-off between niche differentiation (resource partitioning) and absolute fitness. In scenario (4), we examine selection dynamics between genes with higher *R*_0_ but slow mitotic recombination and genes with lower *R*_0_ but fast mitotic recombination. Further details are provided in the section titled “Setup of simulations for the four scenarios”.

### Ups groups of *var* genes

*Var* genes diversify rapidly, primarily through mitotic recombination, but also via meiotic recombination and mutation^24^ . The number of circulating *var* genes in a local host population can reach thousands to tens of thousands^13,14,12^ . Despite this immense diversity, *var* genes are structured into major groups based on their 5’-flanking regions (ups), which regulate gene expression: upsA and upsB/C^16^ . Field repertoires (individual parasite genomes) and isolates (sets of genomes co-infecting the same host) consistently exhibit a stable proportion of these groups across geographical regions and transmission intensities, with approximately 20% upsA and 80% upsB/C variants ^16,27,14,26,12^ .

Ups groups determine many *var* gene features^16,27,14,12^ . Compared with upsB/C genes, upsA genes are more conserved across field repertoires and isolates^12^ and occur at higher frequencies. UpsA genes also form a distinct recombination group, sharing few homologous segments with upsB/C genes^36,37^ . Recombination between the two ups groups is restricted, resulting in a hierarchical recombination structure among *var* genes. Moreover, the two ups groups differ in their mitotic recombination rates: upsB/C genes recombine more frequently with one another than upsA genes^24^ .

These contrasting features have led the hypothesis that upsA and upsB/C genes serve distinct biological roles^25^ . UpsA genes are thought to encode functionally constrained, highly virulent *Pf* EMP1 variants associated with severe disease, whereas upsB/C genes are proposed to adapt for fast recombination to facilitate immune evasion, potentially at the cost of receptor-binding efficiency. However, empirical evidence remains mixed, with inconsistent findings across studies^21,38^ .

Investigating how gene composition within multigene families evolves in individual parasite genomes across transmission gradients can clarify whether these ups groups correspond to distinct transmission strategies. If a stable proportion of genes adapted for each strategy emerges dynamically across transmission gradients, this would support a mapping between transmission strategies and ups categories. Conversely, failure to observe such stability would suggest that ups groups do not map cleanly onto transmission strategies, limiting the ability to identify pathologically important subsets based solely on sequence-based ups classification.

### The stochastic agent-based model of malaria transmission

#### Overview

We extend a previously developed stochastic, continuous-time agent-based model^15^ that explicitly captures feedbacks between population dynamics and parasite evolution. The key assumptions and processes of the model are summarized below and described in detail in the supplementary material, with a schematic overview provided in supplementary figure S2.

Individual human hosts and parasite strains are tracked explicitly. All possible future events–including local transmission, transmission via migrant bites, mitotic and meiotic recombination, mutation, and within-host dynamics such as liver-stage progression and blood-stage progression (which includes the, sequential expression of individual genes within the multigene family of each parasite strain), as well as infection clearance–are stored in a single event queue along with their scheduled execution times. These times may be fixed or drawn from exponential distributions with specified rates. Event rates are parameterized using estimates from the malaria epidemiology literature, related field studies, and in vitro or in vivo measurements. Model parameters, symbols, and values are listed in Supplementary Data 1 SimParams.xlsx. When an event is executed, it may trigger the addition, removal, or modification of future events in the queue, resulting in the recalculation of their scheduled execution times. Simulation are implemented using the next-reaction method^39^, which optimizes the Gillespie first-reaction method^40^ .

The host population size is assumed to be constant: individuals die and are immediately replaced by immunologically naive newborns. Mosquitoes are not modeled explicitly; instead, transmission occurs at an effective contact rate that determines the timing of infectious events. At each event, a donor and recipient host are selected at random. Successful transmission requires active blood-stage infection in the donor and available within-host liver-stage capacity in the recipient. When multiple strains coinfect a host, they compete for limited within-host resources, thereby reducing the transmission probability of each individual strain.

Each parasite strain carries a repertoire of 60 *var* genes encoding variant surface antigens, with each gene composed of two epitopes. During blood-stage infection, genes are expressed sequentially, with only one gene active at any given time. The duration of expression of a gene depends on whether its epitopes have previously been encountered by the host. Hosts acquire epitope-specific immunity upon exposure, and prior immunity shortens the expression time of corresponding genes. Infection terminates once all genes in the repertoire have been expressed and cleared; thus, strain-level infection duration emerges from the number of previously unseen epitopes carried by the repertoire. Finite carrying capacity at both the liver and blood stages constrain multiplicity of infection (MOI), defined as the number of distinct parasite genotypes or strains within a host at a given time), and limit establishment of additional strains.

Antigenic diversity is generated primarily through mitotic recombination, as well as mutation, and can also be introduced via migrant bites. Mitotic recombination occurs within parasite genomes during the asexual blood-stage of infection, exchanging epitope segments between genes and producing novel variants with a specific probability, subject to a recombination load. Meiotic recombination occurs at transmission and generate recombinant novel offspring strains by randomly assembling genes from parental strains. The specific extensions of the model based on Zhan et al^15^ for the present study are summarized in the supplementary material.

#### Setup of simulations for the four tradeoff scenarios

We consider two discrete values for each of the aforementioned traits–a high/fast/long value versus a low/slow/short value (Table 1, Supplementary Data 1 SimParams.xlsx)^41,42,43,24^ . In scenario (1), newly generated genes may be assigned either a high or low transmissibility, long or short durations of expression, and correspondingly high or low absolute fitness (*R*_0_), as well as fast or slow mitotic recombination rates.

In scenario (2), we assume two groups of genes with either a weak or strong trade-off between transmissibility and duration of expression. Under a weak trade-off case, more transmissible genes (group 1) are associated with a slightly shorter duration of expression, yet still achieve a higher *R*_0_ than less transmissible genes (group 2); that is, the transmissibility advantage is only partially offset by the reduced duration of expression. Under a strong trade-off, more transmissible genes (group 1) exhibit substantially shorter duration of expression than less transmissible genes (group 2), such that the two groups have equal *R*_0_; in this case, the transmissibility advantage is fully offset by the disadvantage in duration of expression. In this scenario, gene innovation via mitotic recombination and mutation is turned off.

In scenario (3), we consider a fast-recombining group of genes (group 1) and a slow-recombining group (group 2). Fast-recombining genes tend to be more diverse and less similar in sequences and are therefore more likely to generate non-functional offspring during recombination, resulting in a higher recombination load. Further details on the specification and calculation of recombination load are provided in the supplementary material (see section “Mitotic recombination”). The two groups are assumed to have identical *R*_0_, with equal transmissibility and duration of expression.

In scenario (4), we again assume two groups of genes. Group 1 genes have higher transmissibility and longer duration of expression than group 2 genes, resulting in a higher *R*_0_. However, group 1 genes have lower mitotic recombination rates than group 2 genes.

In scenario (1), mitotic recombination is allowed between any pair of genes. In contrast, in the remaining three scenarios, recombination occurs only between genes within the same group, and genes from different groups do not share alleles or epitopes. Offspring genes inherit their properties from their parental genes. We assume that the expression order of the 60 genes within each parasite genome is random^44^ ; that is, each gene has an equal probability of being expressed early, regardless of its properties.

#### Initial conditions and the maximum transmission intensity considered

We consider three initial conditions in which strains used to initialize transmission contain either predominantly group 1 genes (80% vs. 20%, corresponding to an 8:2 ratio), equal proportions of the two gene groups (50% vs. 50%, or a 5:5 ratio), or predominantly group 2 genes (20% vs. 80%, or a 2:8 ratio).

We increase the transmission intensity log-linearly by adjusting the effective contact rate. The upper bound of this rate is capped to produce epidemiological and diversity metrics comparable to those observed in high-transmission endemic settings^45^ . This approach yields a spectrum of transmission regimes, ranging from low transmission–characterized by near-zero prevalence and MOI–to high transmission, for which simulated epidemiological quantities match those observed in the field. Each simulation is run for 200 years to allow the system to reach a semi-stationary state, after which 15% of the host population is sampled at the end of the high-transmission season.

#### Pairwise type sharing (PTS) between repertoires and isolates

The pairwise type sharing (PTS) index quantifies the extent of shared gene types between any two repertoires or isolates^46^ . It is analogous to the Jaccard and Sørensen indices used in Ecology^47,48^ . Here, we use a directional version of this statistic^49^, defined as

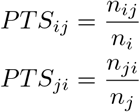

where *n*_*ij*_ denotes the number of gene types shared by isolate *i* and isolate *j*, and *n*_*i*_ and *n*_*j*_ are the total number of unique gene types in isolates *i* and *j*, respectively. This directional formulation reflects that the same number of shared types can represent different degrees of overlap depending on the size of the isolate being considered. The PTS index ranges from 0 to 1, where 0 indicates no shared gene types and 1 indicates complete overlap between the two isolates.

#### The invasion probability of new genes

Below, we summarize the invasion probability of new genes as derived in^15^. The invasion (or fixation) of a new gene^50,51^ occurs when a single copy of a variant present at a given time point leaves descendants in the population after a sufficiently long period. In practice, this requires the variant to surpass a frequency threshold at which it can resist stochastic loss due to genetic drift.

We estimate invasion probabilities using population genetic theory in combination with simulations from our ABM. At stationarity, malaria transmission dynamics can be approximated by a birth-death process under a Moran model with selection. Under this framework, the invasion probability of a rare variant depends on its fitness advantage relative to competing variants^50^, and is given by

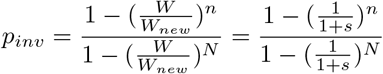

where *n* denotes the number of copies of the new gene, *N* the parasite population size, and *W*_*new*_ and *W* the fitness of the low-frequency gene and that of resident genes, respectively. By convention, resident gene fitness is normalized to 1, and the fitness of the new gene is expressed as 1 + *s*, where *s* denotes its selective advantage.

We assume that the allele pool is sufficiently large that each mutation or mitotic recombination event generates a unique allele, and that back-mutation is vanishingly rare. Consequently, all new variants– whether arising through mutation or mitotic recombination–initially exist as single copies (*n* = 1). Our analysis focuses on parasite populations with effective population sizes *N ≫* 1, under which the expression above simplifies to:

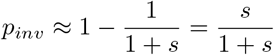

As described by He et al.^52^ for malaria transmission, a gene’s fitness is defined as the expected number of offspring genes it produces, which depends on the population-level transmission rate (*b*), the gene-specific transmissibility (*T*), and the typical infection duration (*d*) of parasites carrying that gene.

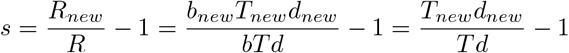

We compare the infection duration of a strain composed entirely of “average” genes with that of a strain containing one novel gene and *g*−1 average genes, where *g* denotes the genome size (i.e., the number of *var* genes per parasite repertoire). An average gene is a theoretical construct whose proportion of susceptible hosts 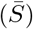 equals the mean susceptibility across all circulating genes. Because the transmission success of a novel gene depends on the total infection duration of the strain in which it is embedded, its invasion probability is influenced by its genomic context (see further below in this section for details).

We derive the expected invasion probability of a novel gene across all possible genome backgrounds, assuming that all other genes in the repertoire are the average genes. Let *d*_1_ and *d*_2_ denote the durations of expression of a gene in naïve hosts, and let *T*_1_ and *T*_2_ denote the corresponding gene-specific transmissibilities. Let *n*_1_ and *n*_2_ represent the average numbers of genes in parasite genomes associated with expression duration *d*_1_ and transmissibility *T*_1_, and with *d*_2_ and *T*_2_, respectively. Since the proportion of susceptible hosts to any novel gene is equal to 1, for a novel gene characterized by *d*_1_ and *T*_1_, we obtain:

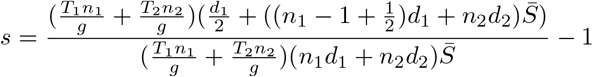

The values of 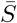 can be computed directly from our ABM simulation outputs.

To further clarify the expression for *s*, we note that because our focus is on the multicopy *var* gene family, the transmission success–or fitness–of a gene depends on the strain in which it is embedded; that is, on the reproductive number of its strain, since transmission occurs at the level of the strain as a whole. Specifically, a strain’s reproductive number–the expected number of secondary infections it generates–is determined by the population-level transmission rate, as well as by its transmissibility and duration of infection, which together govern how frequently a given gene is transmitted to other hosts. Because the population-level transmission rate (*b* for pre-existing and *b*_*new*_ for new genes) is shared across all strains, this term cancels in comparisons of relative fitness.

Strain-level transmissibility (*T*_*new*_ and *T*) is defined as the average transmissibility across all genes constituting a strain. For a strain containing *n*_1_ genes with transmissibility *T*_1_ and *n*_2_ genes with transmissibility *T*_2_, with a total of *g* genes per strain, the mean transmissibility over the course of infection–assuming sequential gene expression–is given by

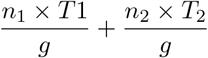

We assume that offspring genes inherit both transmissibility and duration of expression from their parental genes. Consequently, the strain-level transmissibility is identical for a strain composed of one newly generated gene and *g −* 1 pre-existing genes and for a strain composed entirely of pre-existing genes. This term therefore appears in both the numerator and denominator of *s* and cancels.

Differences in fitness instead arise from differences in total infection duration. A strain containing one newly generated gene and *g −* 1 pre-existing genes has a longer duration than a strain composed entirely of pre-existing genes, owing to the prolonged expression of the novel gene. Because only one of the two epitopes of the new gene is truly novel–while the other is inherited from the parental gene–the total infection duration is increased by one half of the duration of expression of a full gene (see the section “Mitotic recombination” in the supplementary material for details).

For preexisting genes and their alleles, the average duration of expression across the host population is scaled by the proportion of susceptible hosts 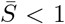, because these genes are expressed only in hosts who are naive to them and are immediately switched off or deactivated in immune hosts. In contrast, newly generated genes–and in particular their novel epitopes–can infect all hosts in the population; thus, the population-level duration of expression of their novel epitopes is not scaled by 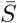. Accordingly, for a strain containing *n*_1_ pre-existing genes with duration *d*_1_ and *n*_2_ pre-exisiting genes with duration *d*_2_, the average infection duration across hosts in the denominators is

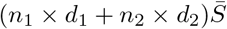

In the numerator for a strain containing one novel gene (one of its two epitopes being novel) and *g* − 1 pre-existing genes, this becomes

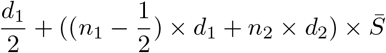

which replaces the term 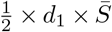 with 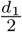, reflecting the prolonged duration of expression of the newly generated gene relative to the pre-existing gene it replaces.

## Results

### No trade-off within or between the two axes (Scenario 1): functional selection optimizes absolute fitness of strains and immune selection increases divergence in *var* gene composition between them

The intensity of immune-mediated competition and immune selection is quantified by the proportion of hosts immune to circulating genes, which increases with transmission intensity (Figure 1a). Under low transmission, competition between co-circulating genes and strains is weak, allowing the coexistence of variants that differ in absolute fitness (*R*_0_) as well as in niche differentiation or resource partitioning capacity, as measured by mitotic recombination rate. Consequently, genes with high transmissibility, prolonged duration of expression, collectively high *R*_0_, and fast mitotic recombination rate do not exclude their inferior counterparts and therefore fail to dominate or become fixed within individual genomes or across the population. This is reflected in mean trait values that remain below their respective optima (Figure 1b-c), as well as relatively high trait variance among circulating genes (Figure 1b-c). In addition, strains and isolates often share a substantial fraction of genes (Figure 1b). These patterns are robust to the initial proportions of genes with different fitness profiles used to seed transmission (see colors in Figure 1).

**Figure 1:**
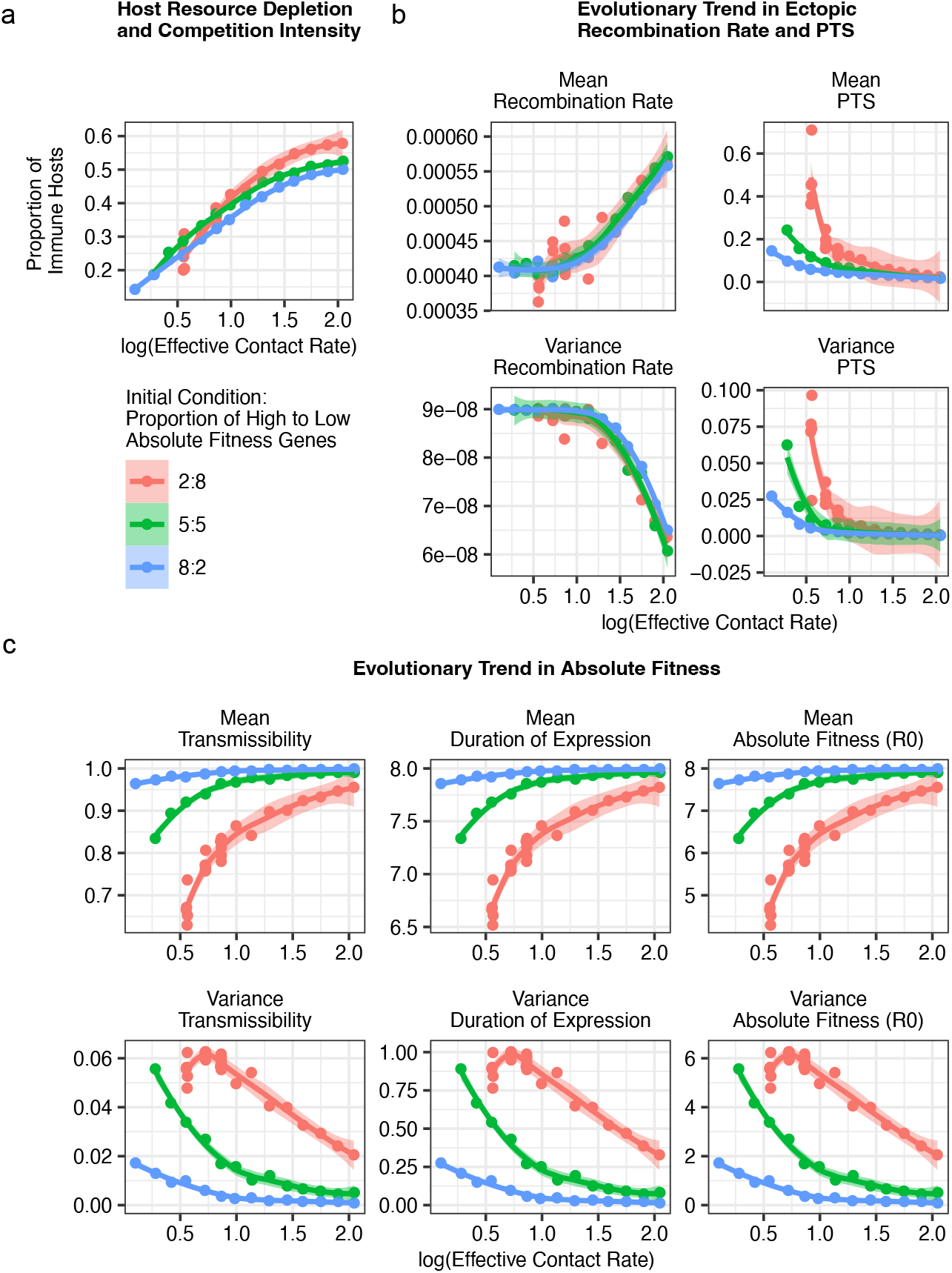
Scenario 1 with no trade-offs. **a** The intensity of immune-mediated competition and immune selection, quantified as the proportion of hosts immune to circulating genes, increases with transmission intensity (effective contact rate; see the section on Transmission intensity in the supplementary material). Under heightened competition, parasite populations evolve towards **b** faster recombination rate and limited gene sharing between strains measured by the pairwise type sharing index or PTS (Methods), and **c** enhanced transmissibility, prolonged duration of expression, and thus higher *R*_0_. Genes with lower transmissibility, shorter duration, or collectively, smaller *R*_0_, and slower recombination rate are progressively eliminated, whereas their superior counterparts are reaching fixation within genomes and the population. Consistently, population-wide variances decrease for **b** mitotic recombination rate and strain gene sharing, and for **c** transmissibility, duration of expression, and *R*_0_. All metrics (y axis) are calculated from end-of-high-season samples in the final simulation year (Year 200). Each point represents an individual simulation run at a specified effective contact rate, with trend lines generated using ggplot2’s geom smooth. stats::loess() is used for less than 1,000 observations; otherwise mgcv::gam() is used with formula = *y ∼ s*(*x*, bs = “cs”) with method = “REML”) function.

In contrast, high transmission intensities competition among co-circulating genes and strains, leading to the elimination of inferior genes and the purging of strains that carry them at high frequency. As a result, parasite populations evolve toward optimal transmissibility, duration of expression, *R*_0_, and recombination rate (Figure 1b-c). In addition, strains and isolates exhibit limited gene sharing, as indicated by a decline in mean PTS with increasing competition (Figure 1b). Consistent with these patterns, trait variance across genes approaches zero (Figure 1b-c). Thus, in the absence of trade-offs, directional functional selection operates independently of immune selection, with both forces becoming stronger as transmission intensity increases.

### A trade-off within the absolute fitness axis (Scenario 2): genes with a high *R*_0_ are favored under intermediate but not high immune-mediated competition

Parasite growth rates and parasitemia levels are governed by the binding affinities of *Pf* EMP1 to host endothelial receptors, as well as other infection-induced within-host environmental factors. Although elevated parasitemia can enhance both transmissibility and virulence, it may also elicit stronger immune responses that accelerate parasite clearance and shorten the duration of expression. We therefore model a trade-off between transmissibility and duration of expression, considering both weak and strong regimes (Methods).

To isolate the effects of trade-offs between traits governing absolute fitness (*R*_0_) on gene competition and persistence within multigene families, we disable gene innovation in the model. Under these conditions, pathogen persistence is sustained through host susceptibility maintained by immunity waning and by the continual replenishment of susceptible individuals via births.

In the weak trade-off case, more transmissible genes are associated with slightly shorter durations of expression and therefore maintain higher *R*_0_ values than less transmissible ones. At low transmission, the system largely preserves the composition of parasite genomes used to initialize transmission. As transmission increases from low to intermediate levels, functional selection favors more transmissible genes with higher *R*_0_, leading to their progressive accumulation within individual genomes and across the population (Figure 2a-b). This regime corresponds to a transmission-limited phase in which parasites have not yet exhausted the host immune space (supplementary figure S4a). Although high-*R*_0_ genes and strains enriched for them are transmitted more frequently, they continue to encounter sufficient numbers of susceptible hosts.

**Figure 2:**
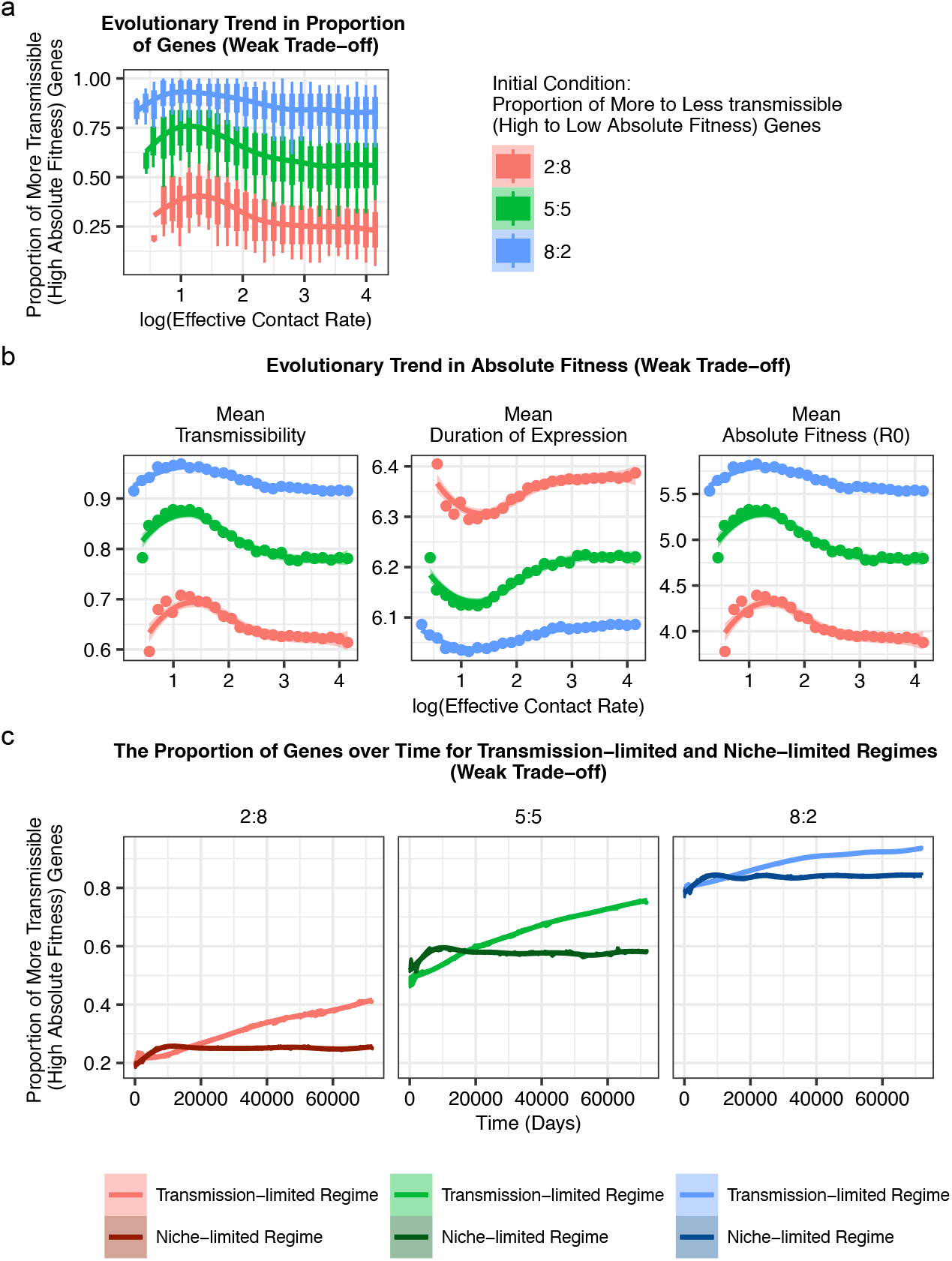
Trade-off between transmissibility and duration of expression. In the weak trade-off case, **a** the proportion of more transmissible (high-*R*_0_) genes initially increases with transmission intensity in the transmission-limited regime, then declines in the niche-limited regime. **b** Correspondingly, population-wide mean transmissibility and *R*_0_ values increase and then decrease, while mean duration of expression shows the inverse trend (initial decrease followed by increase due to the trade-off). **c** Temporal dynamics of the proportion of high-*R*_0_ genes from representative simulations across different regimes reveal distinct trajectories: the proportion steadily rises in the transmission-limited regime (reflecting available immune space) but quickly plateaus in the niche-limited regime (indicating depleted host immune space). Panels correspond to three initial conditions, with color shades matching those in **a**. In **a** and **b**, all metrics on the y axis are calculated from end-of-high-season samples in the last year of simulation (Year 200). Points or boxplots show individual simulation runs with specific effective contact rates. Trends are fitted using ggplot2’s geom smooth function. stats::loess() is used for less than 1,000 observations; otherwise mgcv::gam() is used with formula = *y* ∼ *s*(*x*, bs = “cs”) with method = “REML”) function. In **a**, each boxplot shows the minimum, 5% quantile, median, 95% quantile, and maximum of values across all sampled strains in each simulation.

Conversely, as transmission increases from intermediate to high levels, the proportion of high-*R*_0_ genes declines within individual genomes and across the population (Figure 2a-b). This regime is niche-limited and characterized by intense competition among parasites for hosts (supplementary figure S5a). This reversal arises because high-*R*_0_ genes rapidly deplete available immune space, exposing them to stronger negative immune selection than less transmissible genes. As a result, high-*R*_0_ genes undergo an inversion in selection, shifting from being favored to disfavored as transmission intensity increases.

Beyond examining equilibrium patterns across transmission intensities, we also track the temporal dynamics of the proportion of high-*R*_0_ genes in the population using two simulations representing distinct regimes: one at intermediate transmission (the transmission-limited regime) and one at high transmission (the niche-limited regime), each evaluated across all three initial conditions (Figure 2c). Under intermediate transmission, high-*R*_0_ genes expand continuously throughout the simulation period without reaching saturation. In contrast, under high transmission, high-*R*_0_ genes initially proliferate but rapidly plateau in frequency.

In the strong trade-off case, the increased transmissibility of genes is precisely offset by their shorter duration of expression, resulting in identical *R*_0_ values for more and less transmissible genes. Consequently, both gene groups maintain proportions close to their initial values used to seed transmission (supplementary figure S5b). This scenario effectively represents a neutral system in which no deterministic competitive winner exists, allowing sustained coexistence until stochastic drift ultimately leads to random loss of one gene group.

### A trade-off within the niche difference axis (Scenario 3): recombination load is insufficient to fully counteract immune selection for fast-recombining genes

Although fast-recombining genes generate more novel antigens, they also produce a greater number of non-functional offspring genes as byproducts, resulting in significantly higher recombination load than that of slow-recombining genes (see subsection “Mitotic recombination” under “The stochastic agent-based model of malaria transmission” in the supplementary material, Figure 3a). This recombination load increases with transmission intensity and eventually saturates at high transmission levels (Figure 3a).

**Figure 3:**
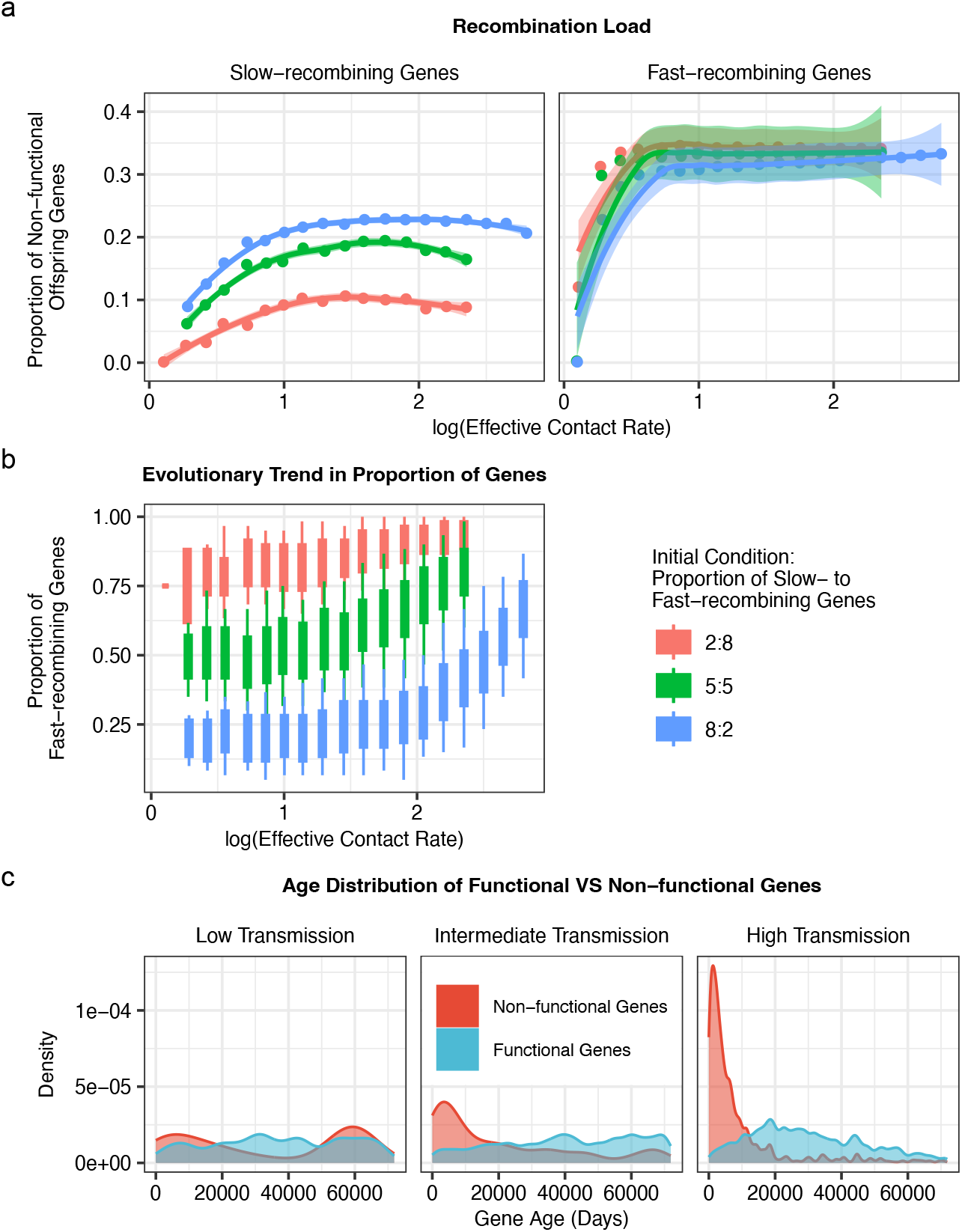
Trade-off between mitotic recombination rate and load associated with mitotic recombination events. **a** Mitotic recombination between fast-recombining genes more often produces non-functional offspring genes than that between slow-recombining genes, generating a higher recombination load. **b** The proportion of fast-recombining genes increases as transmission and immune-mediated competition increases. **c** Age distributions of functional versus non-functional genes using three simulation runs at low, intermediate, and high transmission, respectively. At low transmission, a significant fraction of non-functional genes shows long-term persistence in the population (secondary mode at high ages). At intermediate or high transmission, non-functional genes become more rapidly and efficiently purged (age distribution peaks near zero), while functional genes are long-lived. In **a** and **b**, all metrics on the y axis are calculated from end-of-high-season samples in the last simulation year (Year 200). Points or boxplots represent individual simulation runs with specific effective contact rates. Trends are fitted using ggplot2’s geom smooth function. stats::loess() is used for less than 1,000 observations; otherwise mgcv::gam() is used with formula = *y* ∼ *s*(*x*, bs = “cs”) with method = “REML”) function. In **b**, each boxplot shows the minimum, 5% quantile, median, 95% quantile, and maximum of values across all sampled strains.

At low transmission, the system largely preserves the composition of parasite genomes used to initialize transmission. As transmission increases from low to intermediate levels, functional offspring genes generated through mitotic recombination experience weak selection, because most hosts remain susceptible to existing diversity (supplementary figure S6). As transmission increases further from intermediate to high levels, selection for novel variants intensifies as the proportion of hosts susceptible to existing diversity declines (supplementary figure S6).

Fast-recombining genes are therefore subject to opposing selective forces. Positive selection favors functional novel offspring genes that evade host immunity, prolong infection duration, and enhance transmission of the strains carrying them. In contrast, negative selection removes non-functional variants that shorten infection duration and reduce onward transmission.

Despite this trade-off, the proportion of fast-recombining genes increases within individual genomes and across the population as transmission intensity rises, ultimately approaching fixation (Figure 3b). This indicates that recombination load alone is insufficient to offset the selective advantage of fast-recombining genes under conditions of strong immune-mediated competition for hosts.

The system is therefore resilient to mitotic recombination load at high transmission intensity. To further illustrate this resilience, we examine the age distributions of functional and non-functional offspring genes across the full transmission gradient (Figure 3c). At low transmission, non-functional genes persist for extended periods in the population. At intermediate transmission, non-functional genes exhibit reduced longevity, reflected by the loss of an older-age mode, whereas functional genes show increased persistence, reflected by the loss of a younger-age mode. At high transmission, non-functional genes are rapidly eliminated, approaching zero age, while functional genes predominantly reach intermediate to old ages. Thus, at intermediate to high transmission, the system efficiently purges non-functional offspring genes while retaining functional ones.

What underlies the system’s resilience against recombination load and its efficient purifying selection against non-functional offspring genes? As transmission intensity increases, mixed infections involving multiple strains–arising through superinfection or co-transmission–become increasingly common in highly diverse pathogen systems such as *Plasmodium falciparum* malaria. These mixed infections promote frequent meiotic recombination during the sexual stage in mosquito vectors. Elevated meiotic recombination has two key consequences: enhanced purifying selection and amplified positive selection. By breaking linkage between functional and non-functional genes, which would otherwise constrain adaptation, meiotic recombination brings non-functional genes from different genomic backgrounds together within the same genome, thereby strengthening selection against these genes and the strains that carry them. Additionally, meiotic recombination assembles functional genes into the same genomic backgrounds, increasing both the intensity and efficiency of selection for these genes and their carrier strains.

### Under a trade-off between the two axes (Scenario 4), genes with high *R*_0_/slow recombination rates are selected at intermediate transmission intensity, whereas those with low *R*_0_/fast recombination rates are selected at higher transmission intensity

In scenario 4 (Table 1), we introduce a trade-off between traits governing absolute fitness and those mediating niche differentiation. This configuration creates a tug-of-war between functional selection and immune selection.

As in previous scenarios, at low transmission and small parasite population sizes, the system largely preserves the composition of parasite genomes used to initialize transmission (Figure 4a). As transmission increases to intermediate levels, a substantial proportion of hosts remain susceptible to circulating diversity (supplementary figure S7), indicating limited immune-mediated competition for hosts. Under these conditions, slow-recombining genes with higher *R*_0_ are favored because of their greater transmission success in susceptible hosts, while novel immune-evading epitopes confer little selective advantage. Consequently, selection for fast-recombining genes remains weak.

**Figure 4:**
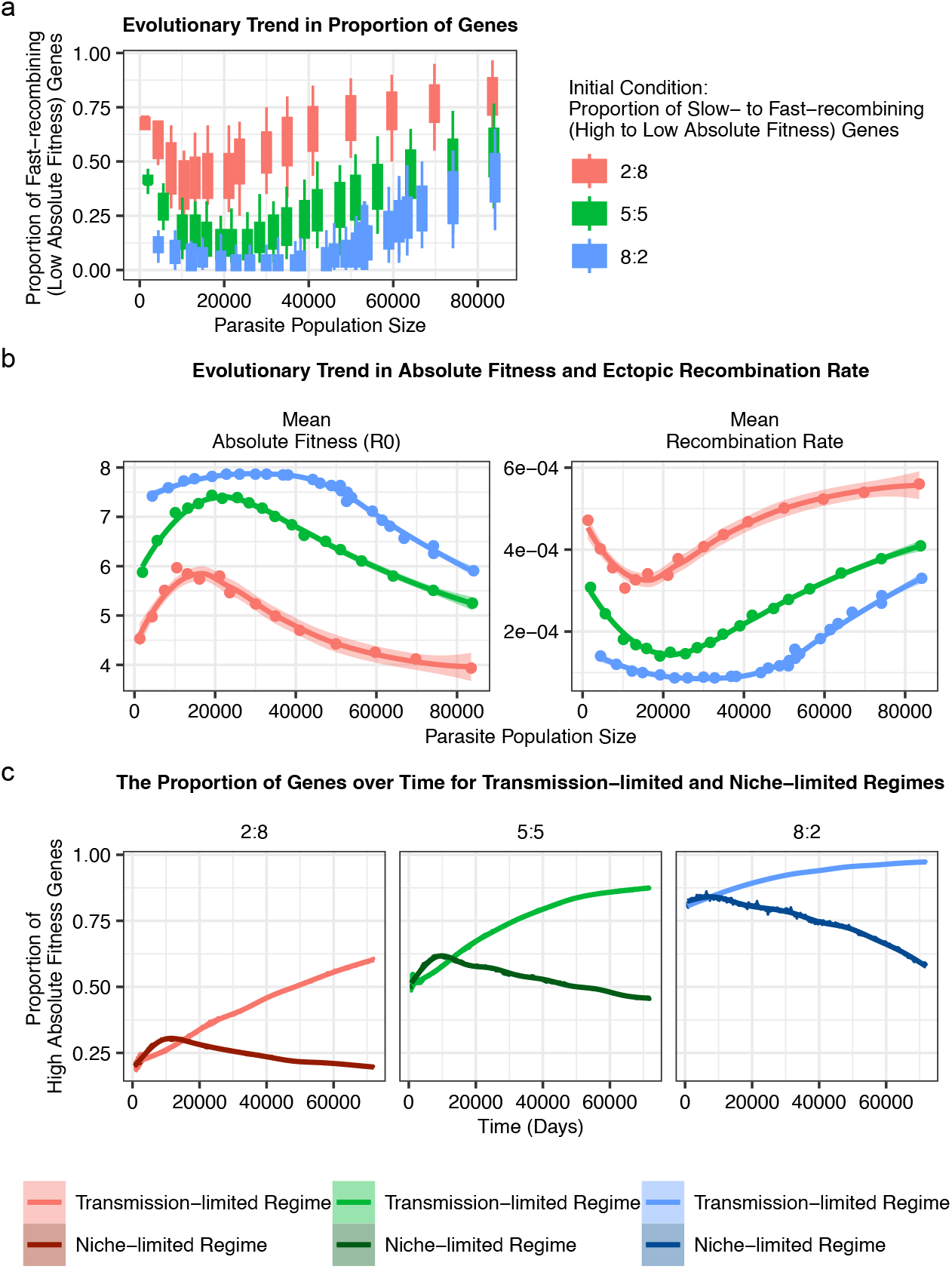
Trade-off between absolute fitness (*R*_0_) and mitotic recombination rate. **a** Slow-recombining (high-*R*_0_) genes are favored at intermediate transmission, but fast-recombining (low-*R*_0_) genes dominate at high transmission. Consistently, **b** population-level mean absolute fitness (averaged across all circulating genes) increases and then decreases with transmission (peaking at intermediate transmission), while mean recombination rate (also averaged across all circulating genes) shows the opposite trend due to the trade-off. **c** Temporal dynamics of the proportion of slow-recombining (high-*R*_0_) genes in the population from representative simulations across different regimes reveal distinct trajectories. In the simulation from the transmission-limited regime, the proportion accumulates progressively in the population. In the simulation from the niche-limited regime, the proportion initially increases and then decreases. The three panels correspond to the three initial conditions with the shades of colors matching those in **a.**In **a** and **b**, all metrics on the y axis are calculated from end-of-high-season samples in the last simulation year (Year 200). Points or boxplots represent individual simulation runs with specific effective contact rates. Trends are fitted using ggplot2’s geom smooth function. stats::loess() is used for less than 1,000 observations; otherwise mgcv::gam() is used with formula = *y* ∼ *s*(*x*, bs = “cs”) with method = “REML”) function. In **a**, each boxplot shows the minimum, 5% quantile, median, 95% quantile, and maximum of values across all sampled strains.

In contrast, at high transmission and large parasite population sizes, host susceptibility to existing diversity declines sharply (supplementary figure S7), reflecting intense competition for hosts. In this regime, slow-recombining, high-*R*_0_ genes encounter transmission bottlenecks due to the depletion of available immune spaces. Gene variants carrying novel, immune-evading epitopes therefore gain a strong selective advantage, and strains enriched for these variants come to dominate transmission (Figure 4a). The resulting decline in the proportion of slow-recombining, high-*R*_0_ genes reflects a strategic shift across the transmission gradient–from a high-*R*_0_, low-mitotic-recombination strategy at intermediate transmission to a low-*R*_0_, high-mitotic-recombination strategy at high transmission (Figure 4b).

Beyond equilibrium patterns, we also examine the temporal dynamics of slow-recombining (high-*R*_0_) gene frequencies, which reveal distinct trajectories across transmission regimes (Figure 4c). In the transmission-limited regime (intermediate transmission), the proportion of slow-recombining genes increases steadily over time as they replace fast-recombining (low-*R*_0_) genes. By contrast, in the niche-limited regime (high transmission), the proportion of slow-recombining genes initially rises but then declines rapidly. This reversal occurs as the depletion of susceptible hosts increasingly favors immune-evading novel variants and, consequently, fast-recombining genes over their high-*R*_0_ counterparts.

### The invasion probability of new genes increases, while the variance of *R*_0_-related traits decreases, with increasing intensity of frequency-dependent selection

Overall, the invasion probability of new genes increases with niche differentiation, quantified as 1 minus the mean PTS among sampled strains, consistent with previous findings under the MCT framework. As niche differentiation increases, both the variance in traits related to absolute fitness and the variance in absolute fitness itself tend to decline (Figure 5, supplementary figure S3-4).

**Figure 5:**
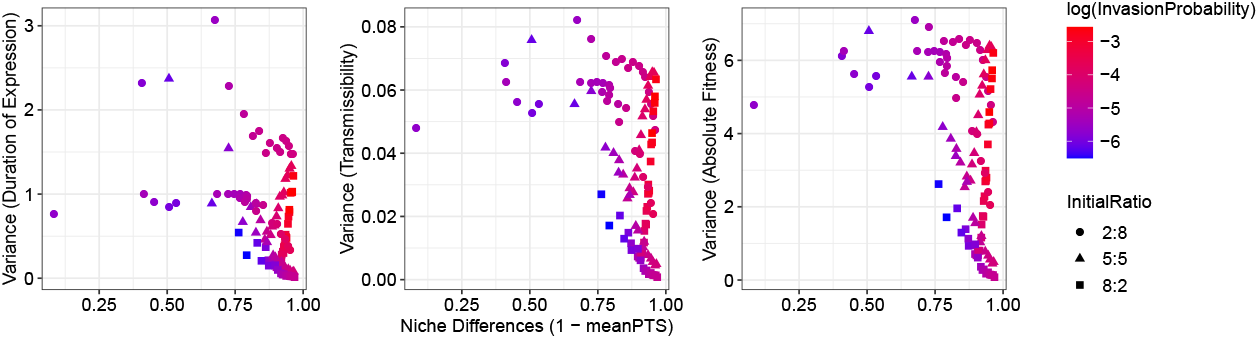
Quantifying the invasion probability of novel genes and the variance in absolute-fitness-related traits and in absolute fitness as niche differences increase. Each point represents a simulation run. The variance of traits related to absolute fitness, including duration of expression and transmissibility, as well as absolute fitness, is calculated across all genes. Niche differences are measured as the divergence in *var* genes between strain pairs, or equivalently as 1 minus the mean PTS across all sampled strain pairs. As niche differences increase, the system shifts towards a state with higher invasion probabilities, where new genes are more likely to invade and become established in the population. Meanwhile, the variance of absolute fitness and absolute fitness-related traits tends to decline. Some runs at high niche differences still show elevated variances due to initial conditions (see Results for details).

With a trade-off between absolute fitness and the mitotic recombination rate, high trait variance can persist, but only as a consequence of the initial conditions–specifically, when transmission is initiated with strains containing equal proportions of high- and low-absolute-fitness genes, or with strains predominantly composed of high-absolute-fitness genes (Figure 4a, c). Each simulation is run for 200 years, sufficient to approach a semi-stationary state and reveal selection trends. Although the system does not fully reach equilibrium and gene proportions continue to evolve, the dynamics exhibit a clear tendency toward fixation of one gene group, accompanied by a progressive decline in variance toward zero (Figure 4c).

The observed pattern of decreasing trait variance with increasing niche differences contrasts with previous results under the MCT framework^53,10^, as discussed below.

## Conclusion

The persistence and turnover of individual member genes within multigene families encoding important surface proteins are shaped by two distinct selective pressures. Directional functional selection favors genes with high absolute fitness, while immune-mediated negative frequency-dependent selection favors genes that enhance antigenic diversification and immune evasion. As parasites evolve to adopt one or both transmission strategies, an enhanced reproductive number vs. immune evasion, trade-offs can potentially operate. To examine long-term evolutionary trends of individual member genes within multigene families, we consider here the *var* multigene family of the human malaria parasite *Plasmodium falciparum*, which encodes the major surface antigen of the blood-stage of infection. The gene composition and gene properties within individual genomes are shaped by the intensity of immune-mediated competition. As expected, low competition allows coexistence of genes with diverse traits, whereas high competition leads to the dominance of genes with optimal traits. We find that when a trade-off exists between absolute fitness and immune evasion, strong competition can weaken directional functional selection for optimal absolute fitness, making differences among all circulating genes more purely frequency-dependent. In this context, traits that promote immune evasion are favored and the proportion of high absolute fitness genes exhibits an inversion pattern across a transmission gradient, shifting from being favored to disfavored as transmission intensity increases. We also find that, regardless of the existence of a trade-off, selection pressures alone cannot precisely maintain a stable proportion of genes adapted to the two transmission strategies within individual genomes across a transmission gradient.

The dynamic variation of the proportion of different groups in our model contrasts with the empirical observation of an approximately constant proportion of upsA and upsB/C genes in field repertoires and isolates across diverse geographical locations and transmission intensities. One possible explanation for this discrepancy is that the ups groups do not map directly onto the two transmission strategies shaped by functional and immune selection. If this is the case, these groups would be unreliable markers of pathogenicity. An alternative explanation is heterogeneity in trait values within each ups group. For instance, some upsA genes may be more prone to mitotic recombination than others, while certain upsB/C genes could exhibit greater virulence. Such within-group variation could help maintain stable proportions of each group in individual repertoires and isolates across transmission intensities, even as selection pressures and the relative abundance of genes adapted to each transmission strategy shift dynamically. Future studies can further test this hypothesis through bioinformatic analyses of sequence data from the two ups groups sampled across geographical locations with varying transmission intensities. Such analyses could assess whether the degree of sequence conservation within each ups group changes with transmission intensity. Additional investigations could aim to infer mitotic recombination rates of upsA and upsBC genes across transmission gradients. Furthermore, association studies linking ups gene groups to disease severity could be refined by incorporating information on local transmission intensity, to determine whether the strength of the association depends on transmission level, specifically, whether at high transmission intensities the association between upsA genes and severe disease symptoms becomes weaker.

The proportion of the two gene groups within individual parasite genomes, respectively adapted to enhanced *R*_0_ and immune evasion, varies with transmission in a way that aligns with biological intuition. As parasites circulate among hosts with varying immune profiles due to different exposure histories, functional and immune selection pressures can change. To transmit effectively across this heterogeneous immunity landscape, parasites benefit from retaining both gene groups. Transmission shapes the proportion of immunologically naïve versus semi-immune hosts in the population, thereby modulating the relative influence of functional versus immune selection.

The inversion pattern in the proportion of high absolute fitness/high virulence genes along a transmission gradient may help explain the paradoxical relationship observed in the field between malaria transmission intensity and severe malaria admissions. Regions with high transmission often show a lower apparent risk of malaria compared to areas with intermediate transmission^54,55^. Similar sign inversions, influenced by population size, have been reported in other systems where trade-offs exist between innovation rates and absolute fitness^56^.

Pathogens with multigene families that encode important surface proteins are resilient to the load associated with gene innovation. A high copy number reduces the dependence of a pathogen’s fitness on individual member genes, thereby relaxing selective constraints. Meiotic recombination between co-infecting strains can disrupt the positive linkage between functional and non-functional genes, allowing functional novel genes to be integrated into the same genomic backgrounds, which enhances positive selection for these genes and their strains. At the same time, meiotic recombination helps purge non-functional genes by bringing them into the same genomic backgrounds where they are more effectively eliminated. Both processes reflect the Hill-Robertson^57^ and the Fisher-Muller effects^58^. Together, gene copy number and meiotic recombination allow pathogens with antigen-encoding multigene families to harness the benefits of accelerated evolutionary innovation while minimizing the potential load incurred in this process.

Our model also does not account for generalized adaptive immunity to more conserved antigens. While such immunity can reduce parasitemia and clinical symptoms (e.g., merozoite opsonization)^59^, it does not necessarily prevent re-infection^26,60^. This is evident in high-transmission regions where asymptomatic infections are highly prevalent and contribute to ongoing transmission^61,62,63^. We therefore assume that infection duration is mainly governed by variant-specific immunity, which could be shortened by the inclusion of generalized immunity. However, we expect that incorporating generalized immunity would not substantially alter the patterns or trends of the evolutionary outcomes identified in our study, as analysis of empirical data shows strong signatures of variant-specific immune selection shaping the diversity and population structure of *var* genes in falciparum malaria^49,26^.

Our study investigates coexistence and diversity in a host-pathogen system with a high-dimensional trait space while explicitly incorporating feedback between evolutionary processes and ecological dynamics. The observed reduction in the variance of traits as niche differences grow, contrasts with previous findings under MCT, where variance in absolute fitness can typically exhibit higher values at higher niche differences. A key distinction is that in our system evolution is explicit and interacts with population dynamics of the disease and the associated competition for hosts. Despite this difference, we find that the invasion probability of new genes increases with niche differences, consistent with previous MCT findings. The “equalizing” of traits associated with (gene) demography would make the system appear ecologically neutral, even though it is under strong selection from specific antigenic differences underlying competition for hosts.

The interplay between directional selection and negative frequency-dependent selection, whether acting on traits encoded by the same or different genomic loci, and with or without trade-offs between them, arises across both pathogen and non-pathogen systems. Among pathogens, examples include *Streptococcus pneumoniae*^64,65^, in which directional selection acts on loci affecting transmission-related traits such as pathogen load and within-host persistence, whereas negative frequency-dependent selection shapes other genomic regions through distinct ecological processes, including human host–pathogen interactions mediated by serotype-specific antigens, bacterium-mobile genetic element interactions, bacterium-bacterium interactions (e.g., via bacteriocins), resource competition, and antibiotic resistance. Another case is the CRISPR-Cas system^66^, an adaptive immune mechanism encoded in the accessory genomes of many bacteria and archaea, where hosts exhibit variation in intrinsic growth rates and immune profiles subject to both directional and frequency-dependent selection. This interplay also manifests in non-pathogen systems such as Batesian mimicry^67^ and species with alternative male mating strategies (e.g., Uta stansburiana lizards or bluegill sunfish)^68,69^. Collectively, these examples illustrate how directional selection and negative frequency-dependent selection jointly shape diversity and evolutionary trajectories across biological systems. In contrast to the *var* multigene family of *Plasmodium falciparum*, where transmission intensity governs the relative importance of the two selective pressures, in these other systems, the balance between the two may shift with ecological or environmental changes, such as fluctuations in the abundance of protected model species in mimicry systems or alterations in habitat and climate. Insights from this work may thus inform our understanding of broader evolutionary contexts.

## Code availability

For information on the simulation code and analysis scripts, please see the GitHub repository at: https://github.com/qzhan321/EcoEvoPathogenMultigeneFamilies.

