## supplementary material for "Evolutionary trade-offs between functional and immune selection shape multigene families in pathogens"

### Electronic supplementary material for Evolutionary trade-offs between functional and immune selection shape multigene families in pathogens

#### 1 The stochastic agent-based model of malaria transmission

##### a Population size

Individual human hosts die and are immediately replaced by newborns with no preexisting immunity. The age structure of the host population follows a truncated exponential distribution with a mean age of 30 years and a maximum age of 80 years.

##### b Transmission intensity

Mosquito vectors are not explicitly represented as agents; instead, we model transmission using an effective contact rate (hereafter referred to as the transmission rate), which under some assumptions is equivalent to vectorial capacity and determines the timing of local transmission events. At each transmission event, one donor and one recipient host are selected at random from the host population. Transmission intensity is influenced by the magnitude of this rate: at high transmission intensity, inter-event times are short and transmission events occur frequently, whereas at low transmission intensity, inter-event times are long and transmission events occur infrequently.

Given a transmission event, successful transmission occurs if the donor host carries active blood-stage infections and the recipient host has not reached its liver-stage carrying capacity. Because parasite strains compete for limited within-host resources, hosts cannot sustain an unlimited number of concurrent infections. The carrying capacity parameter is informed by empirical estimates of the maximum multiplicity of infection observed in high-transmission endemic settings, specifically Bongo District in northern Ghana, where this value is approximately 20 [1]. Further details are provided in the section on Within-host dynamics.

During a transmission event, each parasite strain present in the donor host is transmitted to the mosquito with probability equal to the transmissibility of the currently expressed gene–determined by its ups group–divided by the total number of blood-stage parasite strains carried by the donor host. This formulation captures within-host competition for limited resources, which effectively reduces the contribution of individual strains to transmission. Further details are provided in the section on Meiotic recombination.

#### c Spatial configuration of the transmission system

The transmission system is spatially open, with two host populations explicitly coupled through migration. Transmission arises from two sources: local transmission within the focal population and infection introduced via migrant bites. The local transmission rate, $\lambda$, governs the occurrence of transmission events within the population and corresponds to mosquito bites in which both the donor host and the recipient host are randomly selected from the same population. Further details are provided in the section on Transmission intensity.

Exogenous infections occur at rate $\lambda_{ext}$, representing transmission via migrant bites, in which the recipient host is drawn from the focal population while the donor host is drawn from the other population. This term captures the effects of human movement–and, less frequently, mosquito movement–between populations, which are not explicitly modeled but are incorporated through their contribution to imported transmission.

#### d *Var* repertoire structure

Each parasite carries a repertoire of 60 genes. Each gene is represented as a linear combination of two epitopes (i.e., alleles), following empirical descriptions of the two hypervariable subregions within the *var* tag region amplified from field isolates [2]. Each gene encodes a distinct variant surface antigen expressed during the asexual blood stage of infection.

#### e Mitotic recombination

We model mitotic recombination among *var* genes within the same parasite genome during the asexual stage inside the human host. Mitotic recombination is a major mechanism driving *var* gene diversification and occurs during both the sexual and asexual stages of the parasite life cycle [3]. For simplicity, we restrict recombination to the asexual stage and to genes within the same genome. In each recombination event, two genes are randomly selected from a strain, and a recombination breakpoint is chosen at random. Under normal recombination, alleles are exchanged between the two genes, with a specified probability of generating novel alleles, whereas under gene conversion the second gene remains unchanged. In the current implementation, we assume that all mitotic recombination events proceed via normal recombination rather than gene conversion. Newly recombined genes are viable with a probability that depends on the similarity of their parental genes and the location of the breakpoint [4].

$$P = \rho^{\frac{x(d-x)}{d-1}}$$

Where $\rho$ denotes the recombination tolerance, $d$ represents the genetic distance between the two parental genes, and $x$ represents the genetic distance between the offspring gene and one of its parental genes. Both $d$ and $x$ are expressed in units of amino-acid residues and are quantified based on pairwise epitope differences, which are subsequently translated into amino-acid substitutions. Each var gene consists of two epitopes; thus, the pairwise epitope difference can take values of 0, 1, or 2. We assume that approximately five amino-acid substitutions are sufficient to generate a new epitope, such that the average amino-acid distance between distinct epitopes is five. This assumption is motivated by empirical studies of cross-immunity among viral serotypes. For example, in SARS-CoV-2, one unit of antigenic distance (defined as a one-fold reduction in HI titer) corresponds to approximately a 10% reduction in infectivity [5]. At key epitope sites, a single substitution can result in roughly 1-4 units of antigenic distance.

Accordingly, five substitutions would correspond to 5-20 units of antigenic distance, a magnitude sufficient to effectively eliminate cross-immunity between genotypes. We therefore consider five substitutions to represent a biologically plausible order of magnitude–large enough to ensure minimal cross-immunity, yet not unrealistically extreme.

The probability of producing functional offspring genes increases as the genetic distance between parental genes decreases. Because fast-recombining genes tend to be more diverse and therefore less similar to one another, mitotic recombination events between them are more likely to produce non-functional offspring than recombination between slow-recombining genes. Under the assumption of a recombination load, non-functional offspring genes replace their parental genes. Because these genes do not express and are deactivated immediately, they shorten the infection duration of the strain in which they occur and thereby reduce its fitness.

#### f Meiotic recombination

Meiotic recombination occurs between strains during sexual replication inside the mosquito vector. Because mosquitoes are not explicitly represented in the model, we represent meiotic recombination between parasite genomes at the time of a transmission event. In hosts co-infected by multiple strains, competition for within-host resources and immune-mediated regulation can reduce parasite densities and, consequently, transmissibility. Accordingly, we assume that coinfection reduces the transmission success of each individual strain.

Specifically, when $m > 1$ strains co-infect a donor host, an independent Bernoulli trial is conducted for each strain to determine whether it is transmitted during a contact event. The success probability is given by the strain's transmission probability, which depends on the transmissibility of the currently expressed gene and is scaled by a factor of $\frac{1}{m}$. Thus, co-infection reduces the transmission probability of each strain by a factor of $m$. Previous modeling studies [6] have explored a range of assumptions about the effects of coinfection on transmission, from no effect to reductions similar to those assumed here, with the latter generally considered more realistic.

As a result, only a subset $n \leq m$ of strains is transmitted. In nature, this subset would co-infect a mosquito vector, and we assume that either the same number or a smaller number of strains–subject to the liver-stage carrying capacity of the recipient host–is subsequently transmitted from the mosquito to the recipient host. To incorporate meiotic recombination, each strain transmitted to the recipient host is generated by drawing two parental strains, with replacement, from the $n$ transmitted strains. If the two parental strains are identical, the original strain is transmitted; if they differ, a recombinant strain is produced and transmitted. Accordingly, the probability of transmitting an original versus a recombinant strain are $\frac{1}{n}$ and $1 - \frac{1}{n}$, respectively.

Although associations between physical locations and major *var* gene groups are established, orthologous *var* gene pairs between strains are often difficult to identify. We therefore implement meiotic recombination as a process in which genes are randomly selected from the pooled gene sets of the two parental strains. Given that *var* gene locations can be reshuffled through mitotic recombination and gene conversion, this assumption provides a reasonable and tractable approximation of meiotic recombination. The resulting offspring strains share a fraction of their *var* genes with the two parental strains. Consequently, meiotic recombination generates relatedness and increases similarity between parental and offspring strains.

#### g Within-host dynamics

Each strain is explicitly tracked through its entire life cycle, including the liver stage and asexual blood stage in the human host, as well as the sexual stage in the mosquito. As mosquitoes are not modeled explicitly, we delay the expression of each strain in the recipient host by 7 days to account for the sexual stage, corresponding to the time required for gametocytes to develop into sporozoites within mosquitoes. We impose an additional 7-day delay to represent the liver stage, corresponding to the time required for parasites to be released as merozoites into the bloodstream and invade red blood cells. After this 14-day delay, the asexual blood-stage infection and *var* repertoire expression begin. During repertoire expression stage, the host is considered infectious with the currently active strain, and only these active blood-stage strains are transmissible to other hosts.

Gene expression within the *var* repertoire is sequential, and infection terminates once the entire repertoire has been exhausted. The duration of expression of a given gene reflects the time required for the host immune system to clear the variant surface antigen it encodes. Consistent with empirical evidence that *var* genes are expressed largely sequentially [7, 8], we therefore assume that only a single *var* gene is expressed at any given time.

The duration of expression of an active gene depends on the host's immune history. If the host has previously encountered both epitopes of a given *var* gene and retains immunity to it–subject to waning at empirically estimated rates [9, 10]–the corresponding parasite subpopulation is rapidly cleared. We capture this process by assigning such genes a very high switching rate (1000), corresponding to a short mean duration of expression of $\frac{1}{1000}$ days. Operationally, the time to switch to the next gene (equivalently, the duration of expression of the current gene) is drawn from an exponential distribution with this mean.

In contrast, when the host is immunologically naïve to a given *var* gene, the immune system requires approximately seven days to mount an effective antibody response, leading to a rapid decline or clearance of the expressed variant [11]. This $\sim$ 7-day timescale is consistent with the duration of individual peaks or waves of parasitemia observed in naive *Plasmodium falciparum* infections [6, 12], which typically arise from expression of a single *var* gene and occasionally from a small number of genes. We therefore adopt a range of 6-12 days for the duration of expression of a single gene. Accordingly, in naïve hosts, we model gene expression durations as exponentially distributed with a mean between 6 and 12 days.

If only one epitope has been seen, the duration of expression is reduced by approximately half. Consequently, the duration of expression of a gene is thus proportional to the number of previously unseen epitopes it carries. Upon deactivation, both epitopes of the gene are added to the host's immunity memory, and the next gene in the repertoire becomes immediately active. A strain is cleared from the host once all *var* genes in its repertoire have been depleted. The total duration of infection of a given repertoire is therefore proportional to the number of epitopes not previously encountered by the host, aggregated across all individual *var* genes in that repertoire.

In addition, we impose carrying capacities for both the liver and blood stages of infection. We set the carrying capacity for each stage to 20, corresponding to the maximum multiplicity of infection observed in empirical data [1]. When the number of strains in the liver stage reaches this limit, the host no longer acquires additional infections when selected as a recipient in transmission events. When the number of blood-stage strains reaches the carrying capacity, strains present in the liver stage are prevented from entering the bloodstream and thus fail to transition to the blood stage. In our implementation, such strains are assumed to be lost rather than queued indefinitely.

##### h Major extension of the model from [13]

The major extensions are as follows. Genes from different groups can be assigned distinct transmissibility, duration of expression, and mitotic recombination rates. Mitotic recombination can be specified to occur either exclusively among genes within the same group or between genes from different groups. In addition, genes from different groups can be defined to exhibit either high or low sequence divergence, depending on whether their epitopes are drawn from a shared pool of possible alleles or from disjoint allele pools.

#### 2 Supplementary Figures & Tables

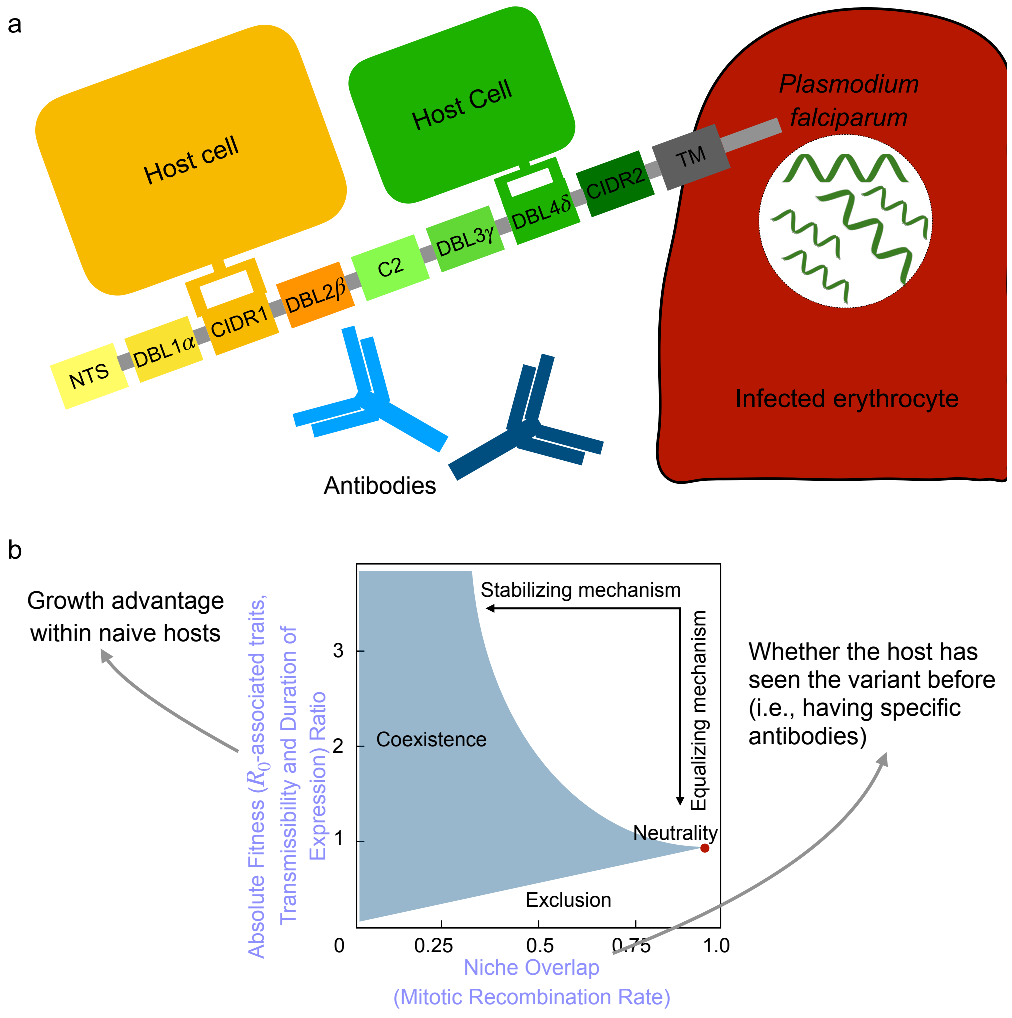


supplementary figure S1: Functional and immune selection of *var* genes and the resulting two different transmission strategies. **a** Schematic representation of a parasite-derived PfEMP1 variant on the surface of an infected erythrocyte. PfEMP1 comprises multiple segments, including an N-terminal segment (NTS), several Duffy Binding-Like (DBL) domains, multiple Cysteine-rich Interdomain Region (CIDR) domains, and a transmembrane domain (TM). Some PfEMP1 variants bind more effectively to host receptors, resulting in enhanced survival or replication in naive hosts. Others are encoded by newly generated genes arising through mitotic recombination, immigration, or mutation, enabling them to evade host immunity established during previous infections. **b** Conceptual framework based on Modern Coexistence Theory (Remade from Song et al [14]). Following the direction of the arrows, stabilizing mechanisms reduce niche overlap, whereas equalizing mechanisms bring the fitness ratio closer to 1. The blue region indicates combinations of niche overlap and fitness ratio compatible with coexistence; the red point represents neutrality. *Var* genes and their PfEMP1 products can adopt one or both of two distinct transmission strategies along two axes: optimizing $R_0$ or promoting rapid innovation for immune escape. Trade-offs [15] may operate along each axis individually, as well as between them.

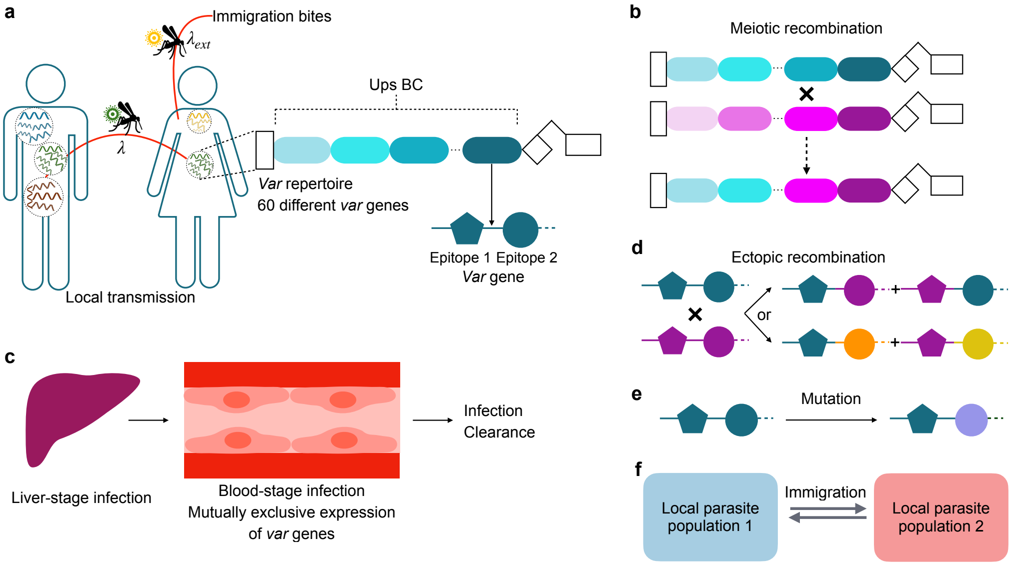


supplementary figure S2: Schematic illustration of the stochastic agent-based model for malaria transmission. **a** Transmission arises from two components: local transmission ($\lambda$) and transmission via migrant bites ($\lambda_{ext}$). Details are provided in the section entitled "Spatial configuration of the transmission system" under "The stochastic agent-based model of malaria transmission". At each transmission event, one donor and one recipient host are selected at random from the host pool. Transmission happens when the donor host carries active blood-stage infections and the recipient host has not reached carrying capacity in its liver. Each parasite genome in the donor host is transmitted to the mosquito with probability of 1/(number of genomes) multiplied by the transmissibility of the currently expressed gene. Each parasite genome consists of a repertoire of 60 *var* genes. Each *var* gene is in turn represented as a linear combination of two epitopes (depicted by different shapes), with each epitope having many possible variants (alleles, depicted by different colors). **b** During the sexual stage of the parasite (within mosquitoes), different parasite genomes can exchange their *var* genes through meiotic recombination to generate novel recombinant repertoires. The recipient host can receive either recombinant genomes or original genomes. **c** When a repertoire is successfully transmitted to a recipient host, it first goes into the liver (dormant) stage during which it is not transmissible. After that the repertoire enters into the blood stage with sequential expression of *var* genes. If the host has no immunity against both epitopes of a *var* gene, the total duration of expression of the gene will last at the order of 10 days (6-12). Immunity against one of the two epitopes will shorten the duration of expression by approximately half. Complete immunity against both epitopes will result in immediate clearance of the gene product. The infection ends when all the *var* genes in the repertoires have been expressed. **d** During the asexual reproduction stage of the parasite, i.e., the blood stage of infection, *var* genes within the same genome can swap their two epitope alleles through mitotic recombination. New epitopes can be generated with a certain probability (see the section on Mitotic recombination for further details). **e** *Var* genes can also mutate their epitopes resulting in new genes. **f** The spatial configuration of the simulated systems is open, and two explicitly coupled local parasite populations are modeled, exchanging migrant genomes with each other.

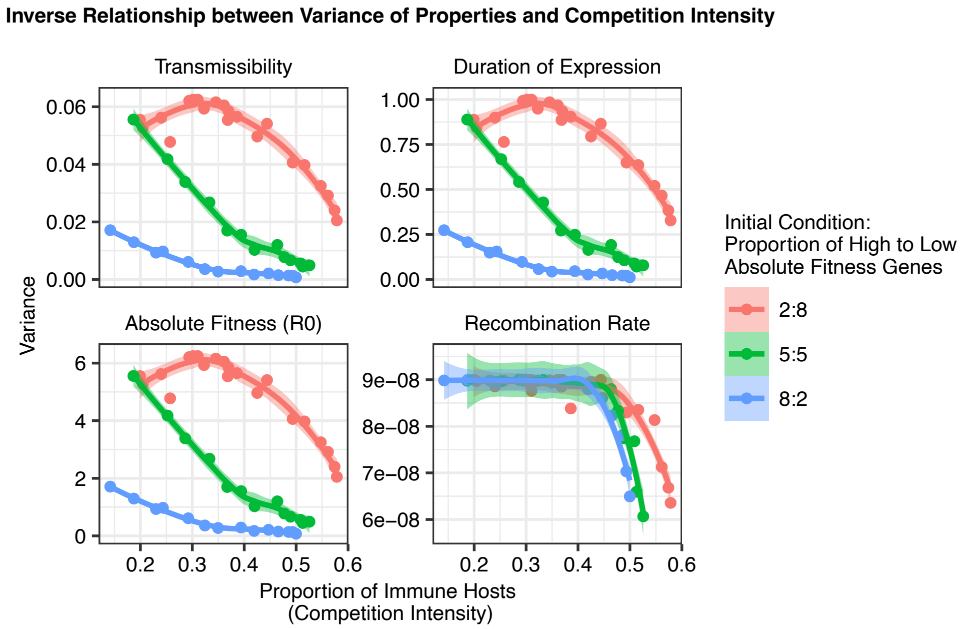


supplementary figure S3: No trade-off within or between the two axes. The inverse relationship between the intensity of immune-mediated competition for hosts and the variance in gene-level traits determining absolute fitness and niche differences, such as transmissibility, duration of expression, absolute fitness itself, mitotic recombination rate. All quantities on the y axis are calculated based on samples taken at the end of the high season in the last year, i.e., 200th year of simulations. Each point represents a single simulation run with a specific effective contact rate, corresponding to a particular level of competition intensity. The trend lines are added using the geom_smooth function in the ggplot2 package. stats::loess() is used for less than 1,000 observations; otherwise mgcv::gam() is used with formula = $y \sim s(x$, bs = "cs") with method = "REML") function.

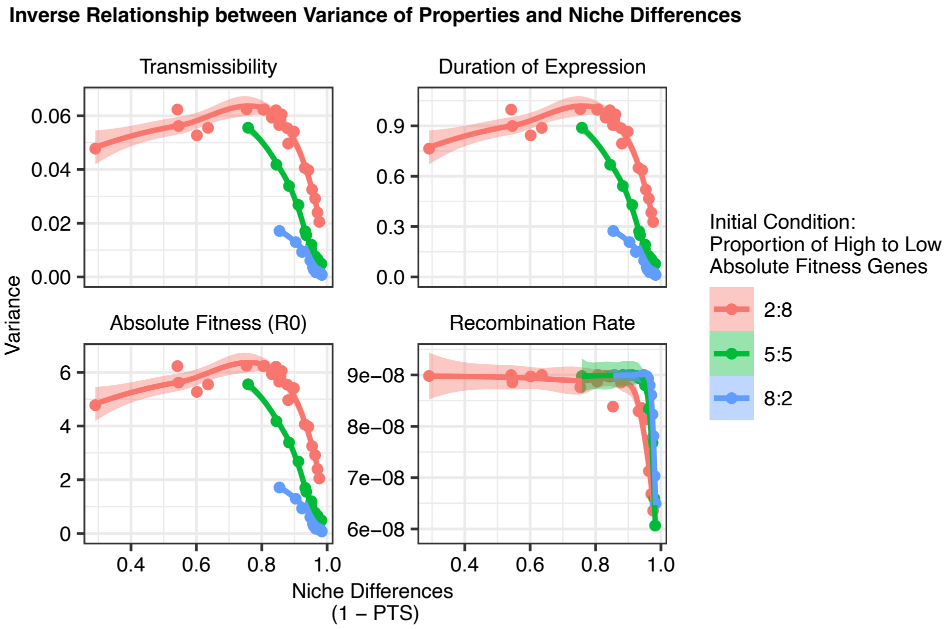


supplementary figure S4: No trade-off within or between the two axes. The inverse relationship between niche differences, quantified as 1− the mean PTS of sampled strains, and the variance in gene-level traits determining absolute fitness and niche differences, such as transmissibility, duration of expression, absolute fitness itself, mitotic recombination rate. All quantities on the y axis are calculated based on samples taken at the end of the high season in the last year, i.e., 200th year of simulations. Each point represents a single simulation run with a specific effective contact rate, corresponding to a specific degree of niche difference. The trend lines are added using the geom_smooth function in the ggplot2 package. stats::loess() is used for less than 1,000 observations; otherwise mgcv::gam() is used with formula = $y \sim s(x, \text{bs} = \text{"cs"})$ with method = "REML") function.

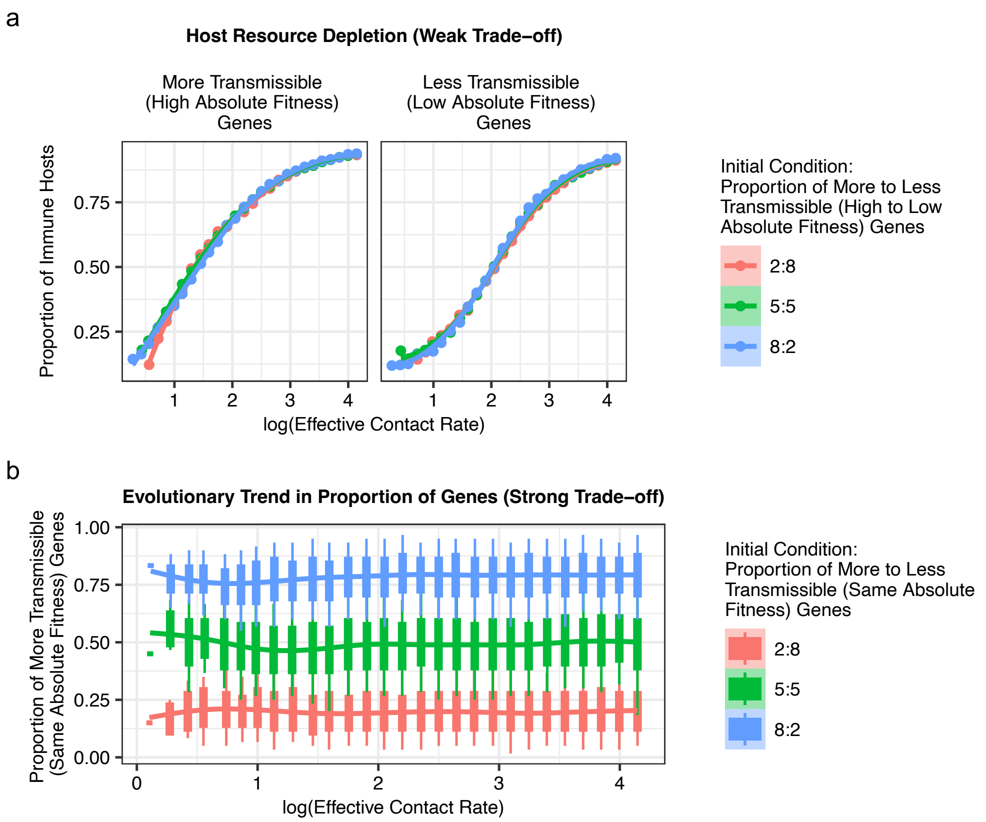


supplementary figure S5: Trade-off in transmissibility and duration of expression. **a** In the weak trade-off case, the intensity of competition, measured by the average proportion of immune hosts to circulating genes, increases with transmission measured by effective contact rate. **b** In the strong trade-off case, the proportion of more transmissible genes remains stable across the transmission gradient. This corresponds to an effectively neutral case in which two groups of genes have the same fitness and thus no deterministic winner exists.

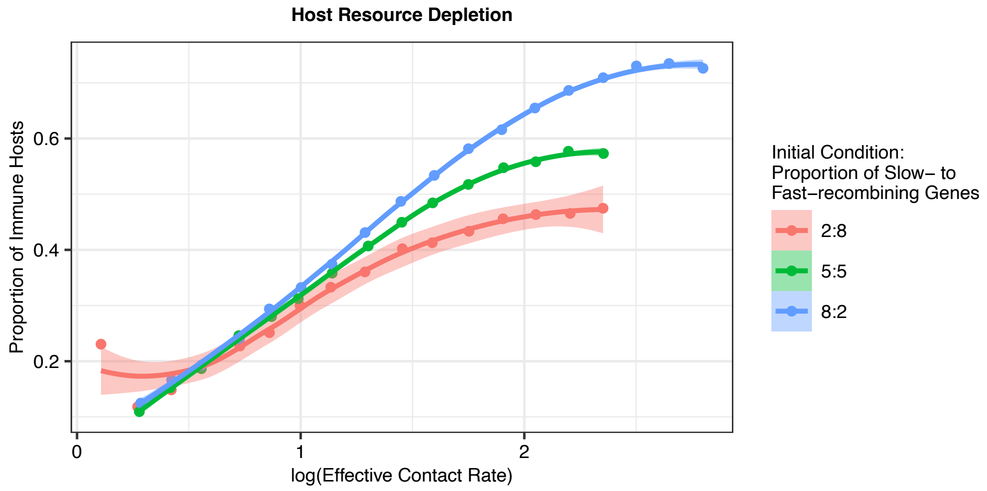


supplementary figure S6: Trade-off between mitotic recombination rate and load associated with mitotic recombination events. The intensity of competition, measured by the average proportion of immune hosts to circulating genes, increases with transmission measured by effective contact rate.

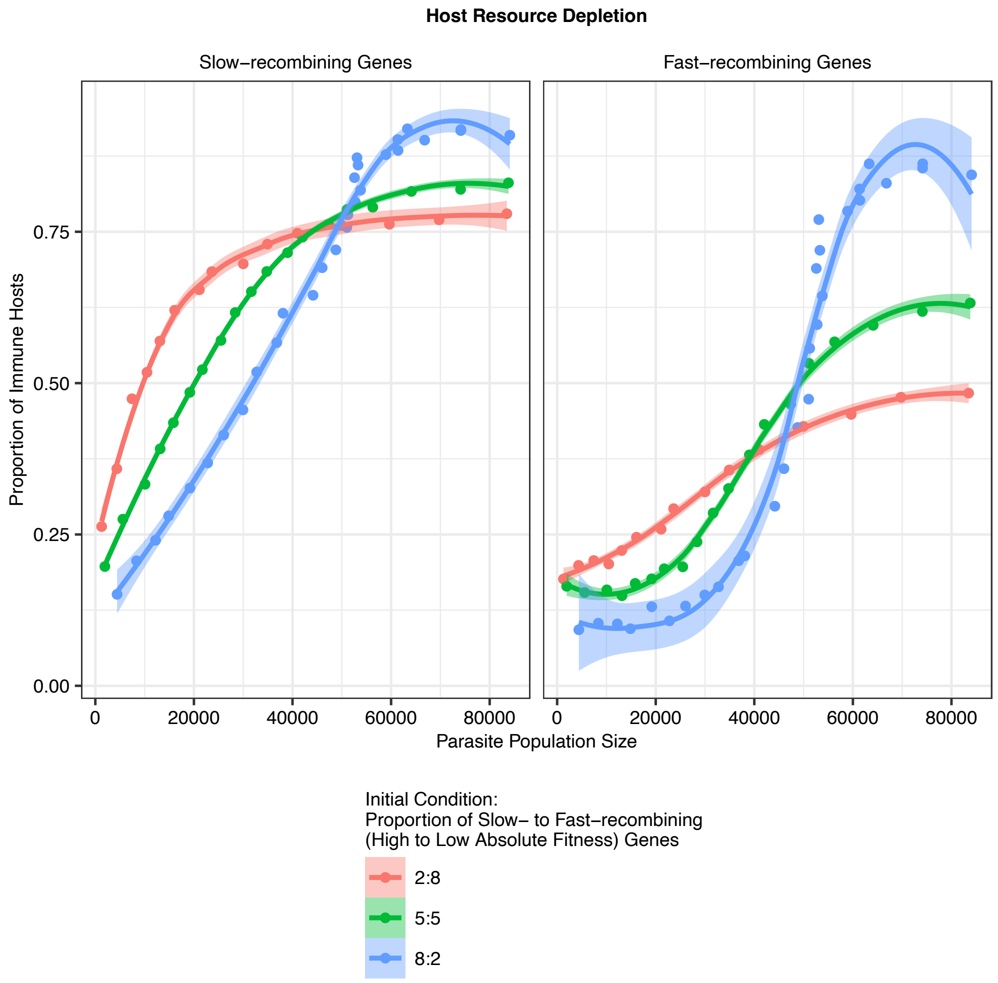


supplementary figure S7: Trade-off between absolute fitness and mitotic recombination rate. The intensity of competition, measured by the average proportion of immune hosts to circulating genes, increases with transmission measured by effective contact rate. Fast-recombining genes generally consume host resource more slowly than slow-recombining genes.
